# Four tumor micro-environmental niches explain a continuum of inter-patient variation in the macroscopic cellular composition of breast tumors

**DOI:** 10.1101/2022.03.04.482793

**Authors:** Anissa El Marrahi, Fabio Lipreri, David Alber, Jean Hausser

**Affiliations:** Department of Cellular & Molecular Biology, Karolinska Institutet & SciLifeLab, Solna, Sweden

## Abstract

The tumor microenvironment is a complex, self-organising tissue whose architecture determines prognostic and response to therapy. There is significant variation in the cellular and spatial architecture of the tumor-microenvironment within and across patients. Are there rules that constrain the architecture of the tumor-microenvironment?

To find out, we develop a quantitative framework of tumor architecture inspired by ideas from satellite imaging, which we apply on deep single-cell profiling and multiplex imaging data in breast tumors. Data analysis shows that inter-patient variation in the macroscopic cellular composition of tumors is structured as a continuum explained by four tumor niches: cancer, inflammatory, tertiary lymphoid structure and fibrotic/necrotic. These niches have their origin in tumor micro-architecture and are shared across patients and tumor subtypes. Niche prevalence depends strongly on the patient, which constraints and explains inter-patient variation.

The present framework will facilitate interpreting inter-patient variation in terms of a tractable number of micro-environmental niches which serve as organizational entities at the meso-scale to bridge the micro- and macro-scales of tumor architecture.

## 1 Introduction

Tumors are complex tissues with a diversity of cell types. Along cancer cells, there are cells of the tumor microenvironment (TME): immune cells, stromal cells, endothelial cells and subtypes thereof [1, 2, 3]. Cells of the tumor microenvironment can represent a sizable fraction of the tumor mass and in some cancer types — lung adenocarcinoma, esophageal adenocarcinoma, breast cancer — they dominate cancer cells in numbers [4].

Contrary to cancer cells, cells of the TME are thought to be mutationally stable. This makes the TME an attractive target for therapy. For example, immune checkpoint therapy interferes with immuno-suppressive receptors of T cells such PD-1 and CTLA-4 in order to boost T cells anti-tumor cytotoxicity [5]. But other cells of the TME can also be targeted in therapy. For example, in glioblastoma, cancer cells secrete signaling molecules such as CSF-1 that cause macrophages to adopt a pro-tumor phenotype [6]. Upon interfering with CSF-1 signaling, macrophages acquire an anti-tumor phenotype, promoting tumor regression [6].

Yet, designing therapies aimed at the TME is challenging because how the multiple cell types in the TME interact to promote or prevent tumor growth is poorly understood. Such interactions have a local nature, taking the form of cell-to-cell signaling through membrane-bound or diffusible signals or metabolic gradients. Unravelling cellular interactions in the TME can thus benefit from spatial methods that map the location and type of cells within tumors, together with signaling molecules and their receptors. Such methods have been developed in recent years, for example in the field of digital pathology [7, 8]. Approaches used in pathology have limitations in terms of the number of RNAs and proteins they can consider simultaneously. To overcome these limitations, approaches based on mass cytometry (MIBI [9], Imaging CyTOF [10, 11]), multiplex immunofluorescence microscopy (t-CycIF [12], 4i [13], Codex [14]) or transcriptomics-based methods (laser capture microdissection [15], MERFISH [16], Spatial Transcriptomics [17], in-situ sequencing [18, 19]) have been developed.

Applying these techniques to tumors has started to reveal the *microscopic* cellular architecture of tumors, at the scale of single cells. A recent study found that colorectal tumors can be partitioned into 9 spatial neighborhoods, each with a stereotypical cellular composition [20]. More generally, a study of 41 in breast cancer tumors found that tumors had either a mixed or compartmentalized micro-architecture, distinguishable on the basis of signaling molecules exhibited by specific immune cells [21].

This spatial micro-architecture of tumors has therapeutic relevance. In the context of immune checkpoint blockade therapy for example, tumors can be stratified into hot, immunosuppressed, excluded or cold based on the density and distribution of T cells within the tumor. Hot tumors have the best response to T cell checkpoint inhibitors, immunosuppressed and excluded tumors show intermediate response and cold tumors respond poorly [22].

Due to the novelty of spatial methods, the potential of spatial approaches for prognosis and therapy is actively being researched. But absent of spatial context, associations between the *macroscopic* cellular composition of tumor samples made of millions of cells and prognosis have been explored systematically. For example, in a majority of cancer types, a higher CD8 T cells content is associated with favorable prognosis whereas a higher macrophage content is worse prognosis, especially if macrophages have an immuno-suppressive phenotype [23].

To reveal how cells of different types and phenotypes coordinate, Wagner et al. [24] characterized the macroscopic composition of 144 breast tumors by mass cytometry. Tumors were found to cluster into 7 types, each with a specific cellular macro-composition [24]. In particular, macrophages expressing the PD-L1 immune ligand and T cells with an exhausted phenotype were abundant in high-grade tumors [24]. Another study of 364 patients across 12 cancer types found 12 types of immune environments, shared across cancer types [25]. Surveys of immunity in more than 10’000 samples from cancer genomics projects found between 4 and 6 immune clusters across 33 cancer types [26] that associate with response to immunotherapy [27]. The immune landscape of tumors has also been characterized through single-cell transcriptomics to reveal the phenotypic heterogeneity of immune cells, phenotypic associations across cell types, and factors controlling this heterogeneity [28].

Much of the thinking regarding the macroscopic and microscopic cellular architecture of tumors has been done in the frame that tumors adopt one out of several possible discrete states. Yet, tumor architecture doesn’t have to be discrete. Many facets of biology are continuous: how tall individuals are, how long they live, the concentration of glucose in the blood. Is there compelling evidence for discrete tumor states or is the cellular architecture of tumors better described by a continuum? Clarifying the discrete or continuous nature of tumor architecture is important to interpreting inter-patient variation in tumor architecture and in informing an appropriate statistical framework to relate tumor architecture to clinical outcomes.

In addition, while tumor architecture has been surveyed both at the macro- and micro-scales, the relation between the macroscopic cellular composition of tumors and its spatial micro-architecture has been little explored. Of macro- and micro-architecture, which of the two has most relevance for prognosis and therapy? Tumor macro- and micro-architecture could be largely uncoupled: much of tumor biology may be encoded in the local micro-architecture of tumors in a way that key information is lost when measuring tumor macro-composition without spatial context. Alternatively, tumor micro-architecture may be strongly constrained by specific rules so that, knowing the macroscopic composition of a tumor, one essentially knows its micro-architecture. How strong is the coupling between tumor macro- and micro-architecture? Clarifying the relation between tumor macro- and micro-architecture can uncover rules governing the interactions between different components of tumors, much like the laws of engineering describe interactions between components parts of a system, a prelude to controlling these components. It could also support clinical thinking on which scale — micro or macro — is most relevant in correlating tumor architecture to prognosis and therapy.

Here, we explore inter-patient variation in the cellular architecture of tumors both at the microscopic and macroscopic scales. We find that inter-patient variation in the macroscopic cellular composition of tumors is a continuum. This continuum is spanned by four archetypal cellular profiles whose contribution to the tumor is patient-dependent. Each archetypal cellular profile represents a tumor microenvironmental niche (TMEN), with four niches explaining spatial variation in the local cellular composition of tumors. Thus, tumor micro-architecture strongly constrains the macroscopic composition of tumors and inter-patient variation. These findings establish a framework that connects the micro- and macro-levels of tumor heterogeneity, to interpret inter-patient variation in the cellular composition of tumors in terms of a small, tractable number of microscopic niches.

## 2 Results

### 2.1 Inter-patient variation in the macroscopic cellular composition of breast tumors is explained by patient-dependent contribution of four archetypal cellular profiles

We begin our exploration of tumor macro-architecture by looking for structure in the inter-patient variation of the cellular composition of tumors. The cellular composition of tumors of 143 primary breast tumors has recently been determined by CyTOF, an antibody-based deep single-cell profiling technology [24]. The abundance of 12 cell types has been measured, so that each tumor can be represented in a 12-dimensional space, where each axis represents a cell type.

Interpreting inter-tumor variation in a 12-dimensional space is challenging: human intuition does not extend beyond 3 dimensions. We thus seek to represent inter-patient variation in tumor cellular composition in a lower dimensional space that accurately captures this variation.

To find this lower-dimensional representation, we perform principal components analysis on the cellular composition of tumors. Principal components analysis looks for directions - or axes - in the 12-dimensional space that spread tumors as much as possible from each other. Looking for axes that spread tumors as much as possible guarantees that the chosen axes best explain inter-tumor variation. In addition, these axes are chosen to be orthogonal to each other, to minimize their redundancy. These axes are called principal components.

We find that cellular composition has a low-dimensional structure: while 12-dimensions are needed to describe inter-patient variation in tumor cellular composition, just 3 principal components already explain 91.7% of the inter-patient variance (Fig. 1A). These 3 principal components represent groups of cell types that vary in coordinated fashion to explain inter-patient variation. Examples of coordination between cell types can be illustrated by plotting the axes of the different cell types on the 2D plane formed by the first two principal components (Fig. 1B). Cell types whose abundance is positively correlated appear as arrows pointing in the same direction - CD4 T cells, CD8 T cells, other immune cells (Fig. 1C). Conversely, arrows pointing in opposite directions represent cell types whose abundance is negatively correlated - immune cells vs cancer cells (Fig. 1D), cancer cells vs healthy tissue cells.

**Figure 1:**
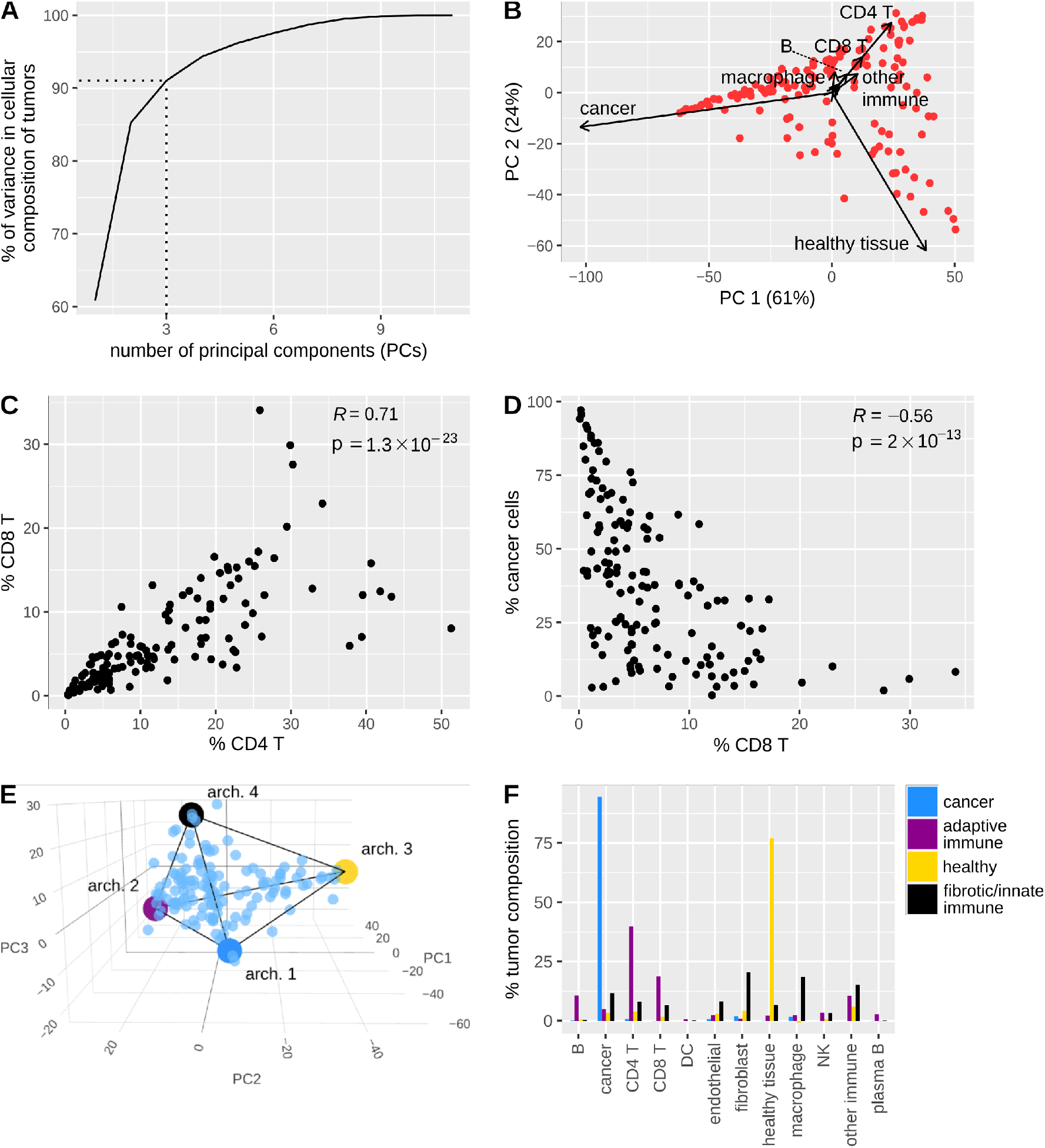
Inter-patient variation in the macroscopic cellular composition of breast tumors is explained by patient-dependent contribution of four archetypal cellular profiles, each with a specific interpretation in terms of histology. **A**. Inter-patient variation in the cellular composition of breast tumors is well described by three axes. **B-D**. There are patterns of correlation and anti-correlation in the abundance of cancer cells, immune cells and healthy cells. Such patterns are revealed by a biplot which project the axes of the different cell types on the first two principal components (B), and by pairwise scatters of cellular abundance (C, D). **E**. Inter-patient variation tumor cellular composition falls on a continuum bounded by a 3D simplex whose vertices represent archetypal cellular composition profiles. **F**. Each archetype has a specific cellular composition and histological interpretation.

Most importantly for the present paper, we find that inter-patient variation is a continuum structured as a 3D simplex, a pyramid with a triangular basis (Fig. 1E). The significance of finding the mathematical structure of a simplex in tumor cellular composition is that any point in a simplex can be written as a weighted average of the vertices - the end points - of the simplex. The vertices of the simplex are computed by archetype analysis, an unbiased, data-driven method from machine learning [29, 30]. These vertices represent archetypical combinations of cells, identical across patients, that are mixed in different proportions depending on the patient. Differences in these mixing proportions explain inter-patient variation in tumor composition (Fig. 1E).

Each archetype of the simplex has a clear interpretation in terms of tissue biology (Fig. 1F). The first archetype represents the cancerous niche, with only cancer cells. The second archetype features mainly CD8 and CD4 T cells as well as B lymphocytes, suggesting an adaptive immune niche. In a third archetype, we find cells from healthy tissue and macrophages. Many juxta-tumoral samples are found close to this archetype (Fig. S1A). This suggests that archetype 3 represents healthy mammary tissue. In the fourth archetype, we find granulocytes (other immune cells) and mesenchymal-like cells, suggesting a fibrotic, innate immune niche. Juxta-tumoral samples are also found close to this archetype.

The low-dimensional simplex is mostly explained by a property of tumor cellular composition: a small number of highly abundant cell types vary most across tumors in an uncoordinated fashion, so that the variance in cellular abundance increases with their mean abundance. Because the sum of cell abundance is 100%, individual tumors land on a simplex where archetypes represent these cell types. Accordingly, shuffling tumor cellular composition to break coordination between cell types while preserving the constraint that cellular composition sums up to 100% captures most of the covariance structure of tumor composition (Fig. S1C).

Yet, not all the covariance structure of the data can be explained by the hypothesis that a small number of highly abundant cell types vary most across tumors in an uncoordinated fashion. There is significantly more structure in the covariance than the shuffling test can explain (Fig. S1C). Under the hypothesis that a small number of highly abundant cell types vary most across tumors in an uncoordinated fashion, archetypes should consist of just one cell type. While the cancer and healthy archetypes verify this, the immune and fibrotic archetypes coordinate multiple cell types. This coordination is also found upon applying a logarithmic transformation to break the relationship between mean and variance, or upon performing PCA on centered-log-ratios [31] to remove the constraint that cell abundance sums up to a 100% (Fig. S1D-L).

While there are known associations between cellular composition and cancer subtype [11], tumors from diverse cancer subtypes are found close to the different archetypes (Fig. S1B). This suggests that the cancer subtype biases but does not absolutely constrains the composition of the tumor [10]. As a result, tumors are made of the same combination of archetypal cellular profiles, shared across breast cancer subtypes.

### 2.2 Four microscopic tumor niches explain the spatial variation in the local cellular composition of triple negative breast tumors

The finding that the macroscopic cellular composition of breast tumors has a recurring architecture across patients constrained by four archetypal cellular profiles bears the question of what is the origin of these archetypal cellular profiles.

The observed archetypal profiles could have a macroscopic or a microscopic origin. In the macroscopic hypothesis, tumors are like a well-mixed solution of cells, with weak local architecture: locally, there is little constraint on how different cell types coordinate and co-localize. Coordination in the abundance of different cell types takes place through long-range signaling interactions and metabolic cues. For example, constraints on the proportions of the different immune cells come about because immune signaling is stereotypical, with different signals attracting different types of immune cells: type 1 cytokines attract CD8 T cells and NK cells, type 2 cytokines attract basophils, type 17 cytokines attract neutrophils and macrophages (Fig. 2A).

**Figure 2:**
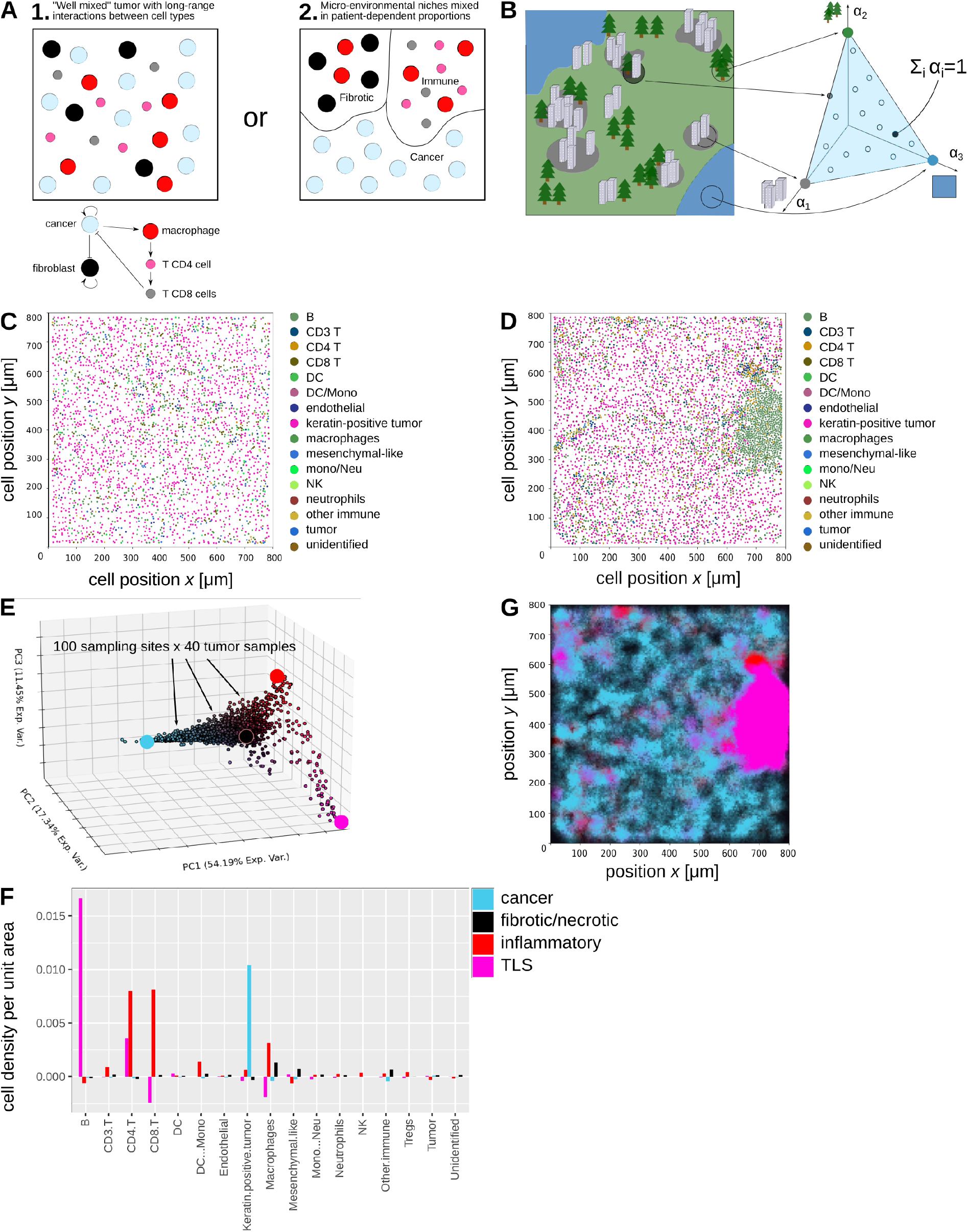
A satellite imaging analogy reveals that four microscopic tumor niches explain the spatial variation in the local cellular composition of triple negative breast tumors. **A**. Two scenarios could explain how inter-patient variation is constrained by archetypal cellular profiles. **B**. If a geographic map can be partitioned into distinct terrain types, local terrain composition must fall on a simplex. Finding the simplex allows discovering the terrain types that compose the geographic map. **C-D**. Cellular maps of two TNBC tumors, from patients 2 (C) and 1 (D). Each dot represents a cell. Colors indicate cell types. Data from Keren et al. [21]. **E**. Local cellular composition across tumors describes a continuum bounded by a 3D simplex in cell composition space, whose vertices represent TMENs. Dots represent the cellular composition of 100 sites of radius from 40 tumor samples **F**. Each TMEN has a specific density in terms of cell types and histology. **G**. Spatial variation in cellular composition is explained by variation in the local prevalence of the four TMENs. The local prevalence of the four TMENs in the cellular map from patient 1 (panel D) is visualized as a color gradient patient.

Alternatively, the observed cellular composition archetypes could find their origin in the microscopic architecture of tumors: different tumors may be built from the same TMENs, each with their own cellular proportions. Inter-patient variation in the prevalence of these niches results in inter-patient variation in the cellular composition of tumors. But because these niches are shared across patients, the macroscopic cellular composition of tumors is constrained (Fig. 2A).

To test this, we note that if tumors have a strong microscopic architecture, one should be able to partition them into the different niches of the micro-environment (or histological areas). This is like partitioning a geographic map into different types of terrains: unbuilt land, urban areas, water bodies, and so on (Fig. 2B). If the map is composed of a finite number of terrain types, the local composition of the map must have the geometry of a simplex. This is because each local region of the map can be broken down into the proportions of the different terrains. These proportions necessarily sum up to 100%, so that any local region of the map can be considered as a weighted average of the different types of terrain - the archetypes of the map.

Just like each tumor niche is composed of specific proportions of cell types, each type of terrain has a specific composition in terms of geographical features: the local number of trees, buildings, water area, and so on. These geographical features must also have the geometry of a simplex: this simplex is connected to the simplex of local terrain composition by a linear transformation that maps each terrain to its geographical features (Fig. 2B, Methods). By analogy, if tumors can be partitioned into micro-environmental niches, the local cellular abundance of tumors must fall on a simplex too.

When the local area considered is large enough, the cellular composition is not predicted to form clusters representing the different micro-environments but rather a continuum within the simplex: this is because sampling local composition randomly over the tumor will consider regions within a given niche as well as regions that overlap between two or more niches (Fig. 2B). This idea, known as hyperspectral unmixing, is used to identify terrain types in satellite images [32].

We test this prediction using a single-cell spatial profiling dataset of 17 cell types in 40 triple-negative breast tumors (Fig. 2C-D, data from Keren et al. [21]). General, recurring principles of tumor architecture are not obvious to visual inspection due to many cell types to consider and strong inter-sample differences (Fig. 2C-D, Fig. S2A).

Despite that, we find that local cellular composition is indeed a continuum bounded by a simplex, as computed by archetype analysis [29]. This simplex lives in a 3-dimensional space whose axes represent the first 3 principal components of local cellular composition (Fig. 2E). Three principal components capture 82% of the spatial variation in cellular composition (Fig. 2E), an observation which is robust to the size of local surveyed area (Fig. S2B heatmap of Gaussian width vs PCs vs %var). Thus, spatial variation in the cellular composition of the tumor is well captured by the simplex.

We find that the archetypes of the simplex have clear interpretation in terms of tumor biology (Fig. 2F). The first archetype represents dense TMENs of cancer cells expressing carcinoma markers such as keratins, p53, and beta-catenin (Fig. S2C). The second archetype features mostly macrophages and mesenchymal-like cells, at low density and could thus represent the fibrotic, necrotic niche (Fig. 2G). In the third archetype, we find a mix of CD68 and MHC-II-expressing macrophages (Fig. S2D marker macrophages), CD8 and CD4 T cells, and NK cells (Fig. 2H). CD4 T cells do not express Foxp3, suggesting a T helper phenotype (Fig. S2E markers CD4). This combination of cell types suggests a type 1 inflammatory response whose function is to trigger anti-cancer immunity: macrophage present antigen on MHC-II to CD4 T cells which then stimulate the cytotoxic activity of CD8 T cells and NK cells [33]. The fourth archetype is characterized by MHC-II and CD45RO expressing B cells and CD4 T helper cells (Fig. S2F-S2G). MHC-II and CD45RO are both expressed by B cells activated by antigen recognition, with B cells presenting the found antigens to CD4 T helper cells for validation [33]. This archetype may thus capture a tertiary lymphoid structure (TLS).

Spatial variation in tumor cellular composition is thus explained by a spatially varying mix of four micro-environmental niches (Fig. 2G, Fig. S2H).

### 2.3 The four microscopic tumor niches explain inter-patient variation in tumor composition

The hypothesis that tumors from different patients can be partitioned into the same TMENs has implications on the structure of inter-patient variation in macroscopic tumor cellular composition: the prediction is that inter-patient variation falls on a simplex in cellular composition space whose vertices represent the four TMENs.

To see why, let 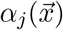 be the local weight of niche *j* at location 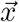 of the tumor. All the weights can be collected into a vector 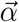, with Σ_*j*_α_*j*_ = 1 and α_*j*_ > 0. We then collect the cellular composition of each niche into a matrix B whose entries b_*ij*_ indicate the density of cell type *i* in niche *j*, in units of inverse volume (1/*μm*^3^). With this notation, the local cellular composition 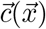 at location 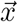 of the tumor is

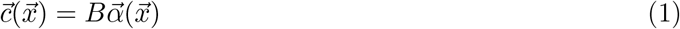

The macroscopic cellular composition of the tumor is then obtained by integrating 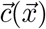 over the tumor volume *V*

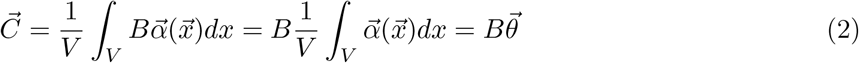

Here, one can show that all θ_*j*_ are positive and sum up to one (Methods). Therefore, the macroscopic cellular composition of the tumor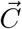 is weighted average of the TMENs B. Macroscopic tumor composition must be bounded by a simplex.

To test this prediction, we compare the macroscopic tumor cellular composition CyTOF data of Wagner et al. [24] with the TMENs found in the MIBI spatial imaging data of Keren et al. [21]. Direct comparison of TMENs and macroscopic cellular composition data is not possible because the tumor samples of Wagner et al. [24] partially include healthy tissue from the tumor margin whereas healthy tissue was not imaged in the samples of Keren et al. [21]. To enable comparison of the two datasets, we mathematically dissect healthy tissue out of the tumor samples (Methods). In addition, the two datasets quantified different types of cells, some of which are shared between the dataset and others are only found in one dataset. We address this by computing cellular composition based on shared cell types (Methods).

As predicted, inter-patient variation in macroscopic tumor cellular composition indeed falls on a low-dimensional simplex bound by the four TMENs (Fig. 3A). The cancer and fibrotic TMENs occupy their own corner of the simplex. But unexpectedly, the TLS and inflammatory TMENs share the same corner despite having different cellular composition.

**Figure 3:**
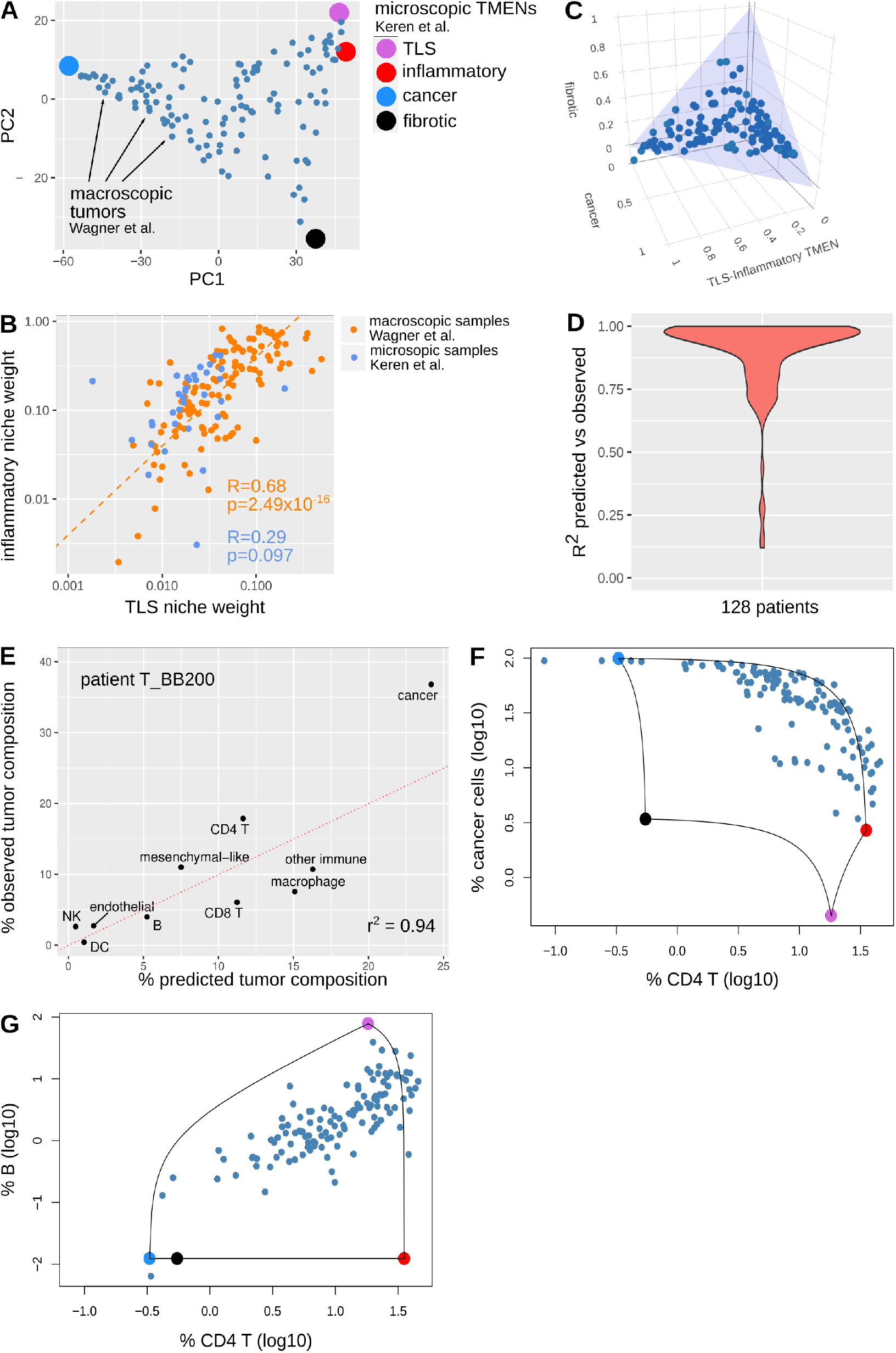
The four microscopic tumor niches explain inter-patient variation in macroscopic tumor composition. **A**. Inter-patient variation in tumor macro-composition is bounded by the four TMENs. The cellular composition of 128 breast tumors (Wagner et al.) was projected on the first principal components after removing the cellular contribution of healthy tissue. The four TMENs computed on the MIBI data from Keren et al. were then projected on the same space. **B**. The inflammatory and TLS TMENs are coupled at the macro-scale but not at the micro-scale. Weight of the inflammatory and TLS TMENs in a regression analysis of tumor composition (dots) on the inflammatory and TLS TMENs. Tumor samples were either macroscopic (orange, Wagner et al.) or microscopic (blue, Keren et al.). **C**. Upon regressing tumor composition on the TMENs, the TMENs weights fall on a unit triangle, consistent with weights summing up to 1. This is expected if TMENs explain inter-patient variation in tumor macro-composition. **D-E**. Tumor macro-composition reconstructed from TMENs highly correlates with experimentally observed macro-composition (median patient in panel E). **F-G**. Inter-patient variation in the pairwise composition of cancer, CD4 and B cells is bounded by the TMENs, over two orders of magnitude.

That the TLS and inflammatory TMENs occupy the same corner of the simplex could be explained if their usage is coupled in tumors. To test this hypothesis, we regress inter-patient variation in the macroscopic cellular composition of tumors on the TLS and inflammatory TMENs. Regression analysis indeed reveals a positive correlation between the weights of the two TMENs (Fig. 3B, *r* = 0.68, *p* = 2.49 × 10^*−*16^). This suggests a macroscopic coupling between the TLS and inflammatory TMENs, with a ratio of 3.5 area units of inflammatory TMEN for each area unit of TLS TMEN. This coupling does not have its origin in microscopic constraints: performing the same analysis on the cellular composition of the 40 microscopic samples of Keren et al. [21], we do not observe a coupling between the TLS and inflammatory TMENs (Fig. 3B, *r* = 0.29, *p* = 0.097).

If patient-dependent usage of the same TMENs explains inter-patient variation in the cellular composition of tumors, one expects that a) regressing tumor composition on the TMENs produces weights that are positive and sum up to 1, and b) tumor composition can be reconstructed based on the TMENs. We find that this is the case. Regression weights fall on a triangular area that represents positive weights and a constraint of summing up to 1 (Fig. 3C, Fig. S3A). Tumor composition is reconstructed from the TMENs with low prediction error (*r*^2^ = 0.95 on average across patients, Fig. 3D-E), including for cell types of low abundance (Fig. S3B). Thus, TMENs accurately model inter-tumor variation in macro-composition.

TMENs constrain the cellular composition of tumors and capture dependencies in tumor macro-composition. TMENs can be used to explain why certain combinations of cells are found in tumors and why others can never be observed. We find that the structure of cellular composition is strongly constrained by tumor micro-architecture. For example, while cellular abundance spans two orders of magnitude in variation, none of the 128 breast tumors of Wagner et al. [24] have high cancer cell content and high CD4 T cell content (Fig. 3F). This is because cancer cells and CD4 T belong to different TMENs and thus their co-occurrence is limited by the areas occupied by these niches. Similarly, there cannot be tumors with many B cells but no CD4 cells: these two cell types occupy the same TLS niche, thus their abundance has to be coupled (Fig. 3G).

## 3 Discussion

The macroscopic cellular composition of tumors is highly individual and of significance for prognosis and response to therapy. Here we ask if there are rules that constrain inter-patient variation in tumor macro-composition. We find that inter-patient variation in the macro-composition of breast tumors forms a continuum is explained by four archetypal cellular profiles, shared across patients and breast cancer subtypes and whose prevalence is patient-specific. These archetypal cellular profiles find their origin in the spatial architecture of tumors which is partitioned in four tumor microenvironmental niches (TMENs): cancer, inflammatory, TLS, fibrotic. These four niches collectively explain most of the spatial variation in the local cellular composition of triple negative breast tumors. Breast tumors from different patients are built from these four TMENs, so that the TMENs explain inter-patient variation in tumor macro-composition.

The observation that TMENs identified from microscopic tumor architecture explain inter-patient variation in cellular composition implies specific rules of tumor architecture: many rules of tumor architecture are incompatible with our observations. For example, if different tumors use different TMENs — analogous to genomics where different tumors have different mutations — microscopic cellular composition does not fall on a single low-dimensional simplex but on more complex shapes. Microsopic tumor cellular composition is not expected to fall on a low-dimensional simplex either if the cellular architecture of tumors is well-mixed (Fig. 2A). If the prevalence of a given TMEN is constant across tumor, microsopic TMENs are not expected to tightly bound inter-patient. Thus, our observations are consistent with the view that tumor architecture is built using four TMENs, shared across patients, and whose prevalences are variable and patient-specific.

TMENs can serve as the foundation for a framework to interpret inter-patient variation in tumor macro-composition and bridge the macro- and micro-scales of tumor architecture. While tumor architecture has mainly been explored, the TMEN framework suggests that tumor macro-architecture may benefit from being considered as a continuum because the prevalence of TMENs in a given tumor is itself a continuum. A benefit of casting tumor macro-composition as a continuum is that this continuum is defined by just 4 archetypes which are more tractable than 7 macroscopic communities or 30 spatial communities previously identified in breast tumors.

TMENs structure and constrain the macro-composition of tumors: many macro-compositions are not allowed if tumors are built from the four TMENs we identified here. Because of this, from measuring the abundance of just 4 cell types of the different TMENs — cancer cells, B cells, T cells, macrophages — one can potentially compute the composition of the tumor in terms of TMENs. Since only a few protein markers are needed for this, determining the composition of the tumor in terms of TMENs is feasible in a clinical setting using established immuno-histochemistry protocols. The strong structure that TMENs impose on tumor composition also opens the door to inferring TMENs and their prevalence in a tumor from non-spatial single-cell data: CyTOF as we illustrate here, but also single-cell RNAseq or flow cytometry data.

Here, we applied an idea from satellite image analysis — hyperspectral unmixing [32] — to identify the TMENs of breast tumors from multiplex images obtained by MIBI. But the same idea can be applied to determine the building blocks of tissues characterized by any method that maps the spatial localization of cells together with their phenotype: other cytometry based imaging methods such as Imaging CyTOF, multiplex fluorescence methods (Codex, CycIF, 4i, and so on) or spatial transcriptomics.

The TMENs we identified correspond to histopathological areas commonly annotated by pathology analysis of tumor sections. Because TMENs emerge independently from analyzing tumor data both at the macroscopic (CyTOF), histological (H&E) scale and at the microscopic scale (MIBI, spatial transcriptomics), TMENs may serve as an organizational principle to structure future efforts aimed at understanding the architecture and dynamics of the tumor micro-environment into 1. researching cell-cell interactions within each TMEN perhaps using notions from tissue homeostasis [34], 2. characterizing interactions between the different TMENs. Finally, in clinical context, TMENs may find use in interpreting inter-patient variability in prognosis in response to therapy.

## 4 Methods

### Processing and analysis of CyTOF data from Wagner et al

We downloaded the summarized version of the CyTOF experiments of Wagner et al. [24]. The data table contains cellular proportions of cell types identified by 73 markers in 144 breast tumors samples. Cellular composition was organized in hierarchies, for example: proportion of live cells among all cells, proportion of cells of the M (myeloid) cluster among live cells, proportion of M1 cells among the M cluster, and so on. Wagner et al. [24] assigned cell types — Tumor Associated Macrophages, CD4+ T regulatory cells, … — to cell clusters (leaf nodes) — M01, T01, and so on. We took over these cellular assignments from Fig. 2(D-L) of Wagner et al. [24] in order to computed the relative composition of each tumor in terms of 12 cell types, chosen to be as similar as possible to the cell types profiled by Keren et al. [21]: cancer cells, fibroblasts, endothelial cells, CD4 T cells, CD8 T cells, NK cells, dendritic cells, macrophages, B cells, plasma B cells, healthy tissue, other immune cells.

One sample contained less than 50% live cells and was thus removed, keeping 143 samples for the analysis.

We performed PCA, centered and unscaled using the ade4 package of the data analysis software R [35].

### Randomization of cellular abundance

To reveal patterns of covariance in cellular composition data not due to coordination between cell types, we shuffled cells proportions cell type-wise. This conserves the overall, marginal distribution of cellular abundance of each cell type. But it breaks the constrain that cellular abundance sums up to 100% in each sample. We thus re-impose this constrain by re-normalizing cell abundance in each sample to 100 %. We performed PCA to collect the fraction of variance explained. We repeated this 100 times to determine error bars on the distribution of variance explained across principal components, that is on the eigenvalues of the covariance matrix.

### Archetype analysis

Archetype analysis [29] aims to fit a d-dimensional simplex as tightly as possible to n data points 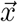. The simplex has *p* vertices 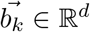, *k* = 1, …, *p* which represent the archetypes. By definition, each point 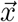 within the simplex can be written as a weighted average of the archetypes

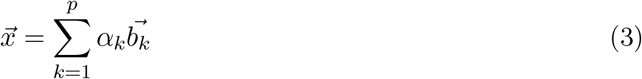

with the weights *α* constrained by 0 ≤ *α*_*k*_ ≤ 1 and 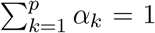. We used the tumor samples projected onto the 3 first PCs as input for archetype analysis. We used the Archetypal Analysis python package [36], with the parameters: n_archetypes = 4, tolerance = 0.001, max_iter = 200, random_state = 0, C = 0.0001, initialize = ‘random’, redundancy_try = 30. The output of this algorithm contains a dataset of α_*k*_ weights for each tumor sample and the coordinates of the archetypes 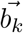 in the reduced space of 3 PCs.

#### Processing and analysis of MIBI data from Keren et al. [21]

From the website of the Angelo lab, we obtained processed MIBI data for 36 protein markers from 41 TNBC patients samples: expression values, segmented images, patient data. In particular, the segmented data (cellData.csv) contained (x, y) coordinates of each cell and its type (out of 17 cell types) as determined by the authors of the study. Following the authors of Keren et al. [21], patient 30 was excluded from the analysis.

#### Measuring density in micro-environmental sites: Gaussian kernel

Each slide is a 2 dimensional space with cells of a determined cell type as points of coordinates (*x, y*). In each slide, we randomly chose 100 micro-environmental sites, by drawing from a uniform distribution across the slide. Each site was defined as the area of width *r* around the random chosen center 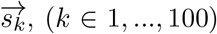. We explored a broad range of width *r* values to examine the robustness of tumor architecture (Fig. S2S). We found that *r* = 25*μm* is the minimal width allowing to capture cellular architecture (Fig. S2B). This suggests that tumor micro-architecture emerges on a scale of 2-4 cells.

To quantify the different cell types in each site, rather than counting cells with the circle of radius *r*, we used a Gaussian kernel density estimation to decrease counting noise: if a cell is slightly outside the circle of radius *r*, it is not counting with the first strategy. Counts are thus sensitive to slight changes in the position of the center of the circle. Gaussian kernel density estimation addresses by weighting cell counts by their distance to the center in a smooth fashion. The weight *g* of a cell of position 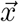 to the site of center 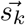 is defined as

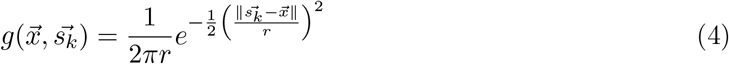

We summed up the density values for each cell type. In total, we generated 4000 sites (in a total of 40 slides).

#### Mapping cells types across two datasets

For the purpose of Fig. 3 and S3, we needed to compare cellular abundances across these two datasets. But the cell types profiled in the CyTOF data of Wagner et al. [24] and MIBI data of Keren et al. [21] overlap only partially: while some cell types are common to both data sets — CD8 T cells for example — other cell types were either only profiled in one dataset, or profiled at a different level of specificity — all B cells vs distinguishing B and plasma cells, all CD4 cells vs distinguishing CD4 cells and Tregs. We thus mapped cell types across the two datasets to compute a set of cell types common to both datasets. To do so, we created an incidence matrix *G* of dimensions *m* × *n* with *m* cell types from MIBI data as rows and *n* cell types from CyTOF as columns. On each cell of the matrix, we put 0 value if two cell types from both data were not similar, 1 otherwise. A column or a row can have more than one 1 if the MIBI and CyTOF datasets differ in their granularity for the corresponding cell type. This incidence matrix can be represented as a bipartite graph (Fig. S3C).

From *G*, we then derive two matrices *G*_*k*_, *G*_*w*_ that allow to project cell proportions from the initial MIBI *X*_*k*_ and CyTOF data *X*_*k*_ (respectively) onto the shared sets of cell types *Y*_*k*_ = *X*_*k*_*G*_*k*_ and *Y*_*w*_ = *X*_*w*_*G*_*w*_ by a matrix multiplication.

To compute the projection matrix *G*_*k*_ for the MIBI data, we initialize *G*_*k*_ := *G*. We then summed up all the rows of G. Rows where the sum is larger than 1 represent MIBI cell types that map to multiple, more granular CyTOF cell types. For these rows, we keep all columns with 0s and the first column with a 1 in order to keep only the least granular cell type of the two datasets. The kept column is then named accordingly to the row. To compute the projection matrix G_*w*_, we perform the same procedure, reversing rows and columns.

Finally the new shared cell types based on which we compare the MIBI and CyTOF datasets were:

| shared cell type | cell type from CyTOF data | cell type from MIBI data |
| --- | --- | --- |
| DC | DC | DC |
| endothelial | EndothelialCells | Endothelial |
| macrophages | Macrophages | Macrophages |
| mesenchymal-like | Other<br>Fibroblasts | Mesenchymal-like<br>Unidentified |
| CD4 T | CD4+T | CD4+T<br>Tregs |
| cancer | EpithelialCells | Tumor<br>Keratin-positive tumor |
| NK | NK | NK |
| B | BCells<br>PlasmaCells | B |
| other immune | OtherImmune | Neutrophils<br>Mono/Neu<br>CD3+T/CD4-T<br>DC/Mono<br>OtherImmune |
| CD8 T | CD8+T | CD8+T |

#### In silico dissection of healthy tissue

Archetype analysis suggested that tumors analyzed by the CyTOF Wagner et al. [24] contained a portion of healthy tissue. To compare these tumors to the MIBI data of Keren et al. [21] which did not profile healthy tissue, the contribution of healthy tissue needs to be dissected out of the CyTOF data.

After projection onto *n* − 1 principal components 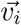, a CyTOF tumor sample 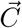 (vector of proportions of 12 shared cell types) can be written as

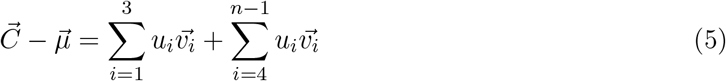

Where 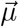 is the vector of average proportions of each cell type and *u*_*i*_ represents the contribution of PC *i* to the sample *C*. We then rewrite the first term as a weighted average of four archetypes (cancer, immunity, healthy, fibrotic) computed by archetype analysis in the space of the first 3 PCs to obtain

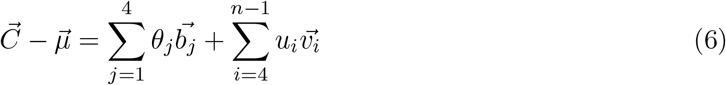

where 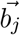 is the *j*th archetype. When then dissect the healthy archetype (archetype 3) by removing its contribution from the weighted averages:

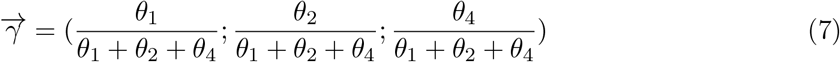

Finally, we computed cellular proportions 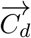 after removing the healthy archetype as

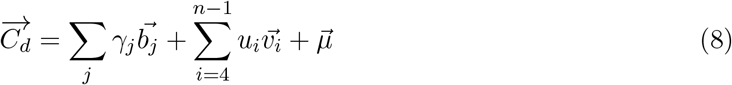

#### Showing that macroscopic TMEN weights are positive and sum-up to one

Here we show that, if macroscopic TMEN weights 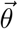 are determined by integrating microscopic cellular composition over the full tumor volume, they must be positive and sum-up to one.

The macroscopic cellular composition of a tumor 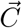 is obtained by integrating microscopic cellular composition 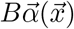 at tumor positions 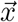 over the tumor volume *V*

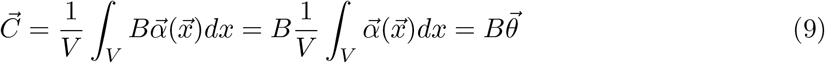

First, since *α*_*i*_ ≥ 0,

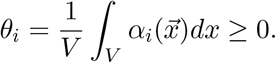

Second,

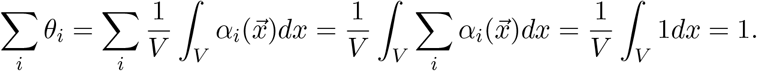

#### Linear regression on macroscopic cellular abundance of tumors

To test if the macroscopic cellular composition of tumors can be modeled by the TMENs, we perform a regression analysis. Let 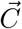 be the vector of cellular composition of one tumor and 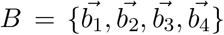 a matrix of cells proportions for the four TMENs, with TMENs as column-vectors 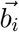. If the cell composition of tumors 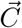 is constrained by a mix of TMENs, it can written as

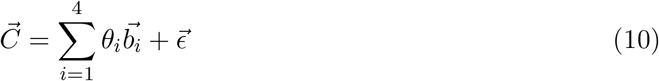

where *θ*_*i*_ represents the proportion of each TMEN in the tumor, 0 ≤ *θ*_*i*_ ≤ 1. 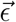 represents the error. With a dataset of many tumors, we can write this equation in a matrix form:

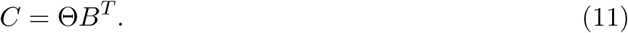

Here, *B* has dimensions of cell types x TMENs. Θ, with dimensions of samples x TMENs, is the unknown in this equation. We solve for Θ using the pseudo-inverse method. To do so, we first multiply both sides by B^*T*^ to obtain

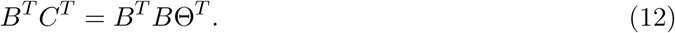

And we solve for Θ,

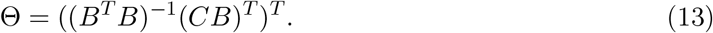

Since (*B*^*T*^ *B*) is a square matrix, we can find its inverse if *det*(*B*^*T*^ *B*) ≠ 0. We numerically found the inverse of *B*^*T*^ *B* using the solve() function in data analysis software R.

We assessed the accuracy of the linear regression by computing the coefficient of determination 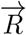 for each sample,

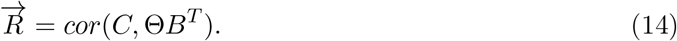

*R*_*i*_ represents the correlation between the cell proportions observed and predicted from a linear combination of TMENs in patient *i*.

## 5 Acknowledgments

The authors thank Petter Säterskog, Antony Cougnoux, Louise Gsell, Guilhem Panneau, Leeat Keren and Benjamin Towbin for discussions and input on this project.

This work was supported by the Swedish Cancer Fund, the Swedish Research Council, SciLifeLab and Karolinska Institutet.

## Notes

### Competing Interest Statement

The authors have declared no competing interest.

