## Supplementary material for "Four tumor micro-environmental niches explain a continuum of inter-patient variation in the macroscopic cellular composition of breast tumors"

March 3, 2022

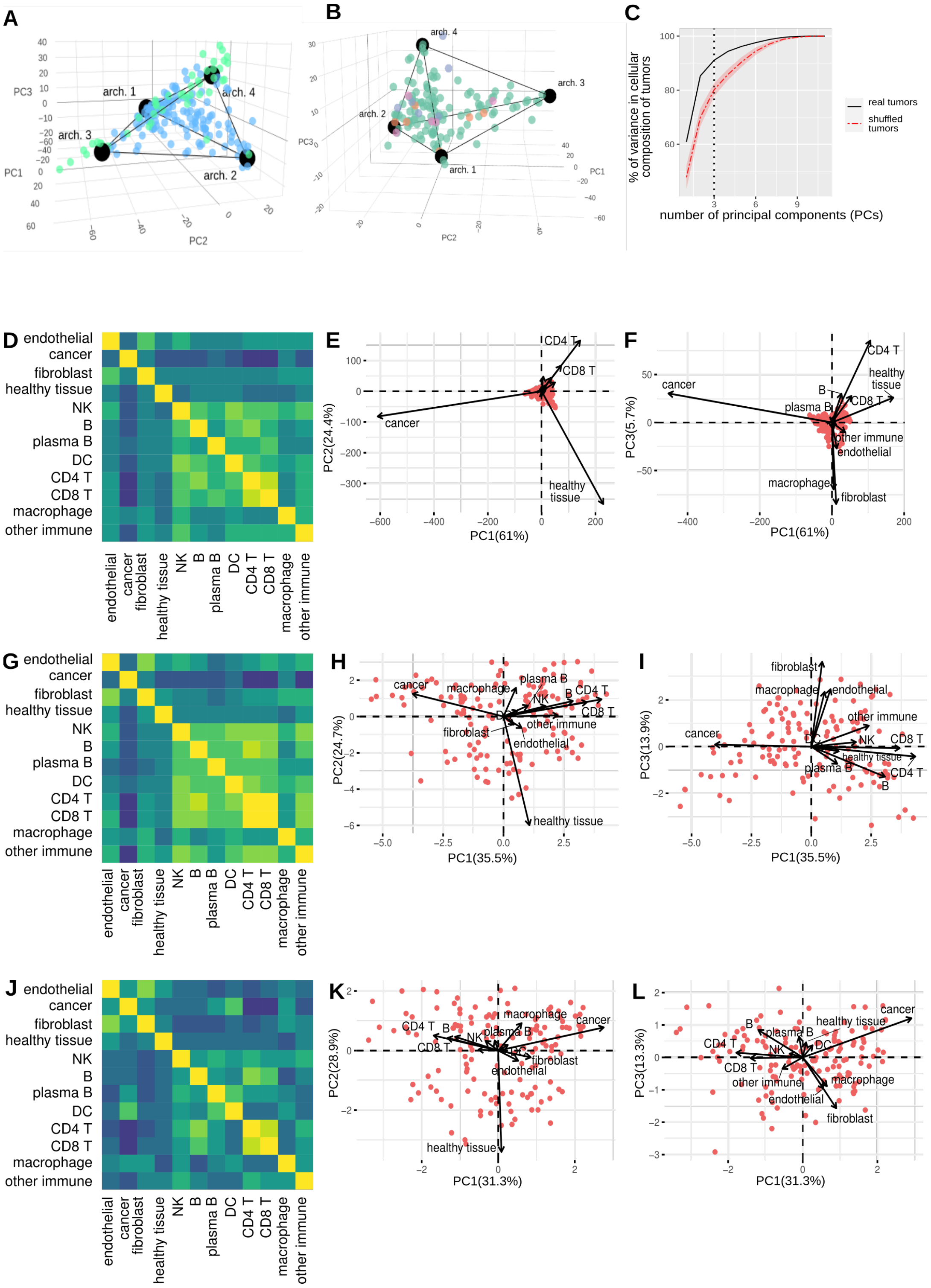


**Supplementary Figure 1: A.** Juxta-tumoral samples (colored in green) map with the healthy and fibrotic archetypes in macroscopic cellular abundance of tumors (colored in light blue). **B.** Sub-classification of breast tumors by hormone receptor response fails to explain inter-patient variability in macroscopic cellular abundance. **C.** A shuffling test that destroys coordination between cell types suggests that much of the structure in the cellar composition of tumors is explained by the fact that 1. a small number of highly abundant cells also vary most in their abundance across patients, and 2. that total cellular abundance is constrained to 100%. Yet, the shuffling test cannot reproduce the full structure of macroscopic cellular composition. This suggests that is significant coordination between the abundance of the different cell types. **D.** Correlation matrix of cell abundances (percentages). **E-F.** Biplots of PC1 vs PC2 and PC1 vs PC3 with cell types (arrows) and tumor samples (red dots) computed on cell abundances (percentages). Samples fall on a tetrahedral structure whose vertices are made of 1. cancer cells, 2. healthy cells, 3. immune cells, and 4. macrophages and fibroblasts. **G-I.** Same for $\log_2$ cellular abundance. Multi-cellular structures are similar to the structures identified when considering unprocessed cellular abundance (D-F). **J-L.** Same for log-centered-ratios. Multi-cellular structures are similar to the structures identified when considering unprocessed cellular abundance (D-F). Data: Wagner et al. [1]

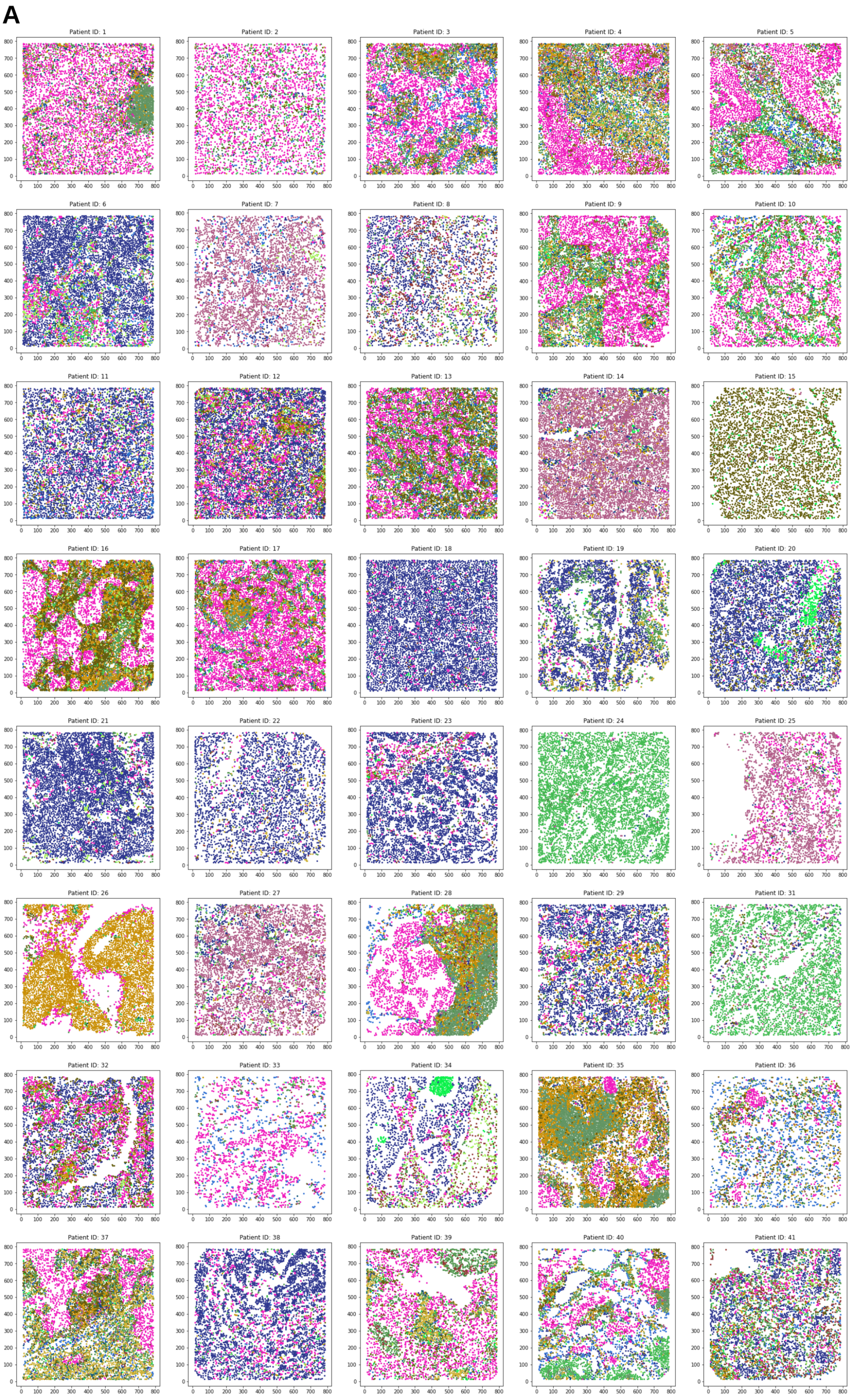
A
Patient ID: 1
Patient ID: 2
Patient ID: 3
Patient ID: 4
Patient ID: 5
Patient ID: 6
Patient ID: 7
Patient ID: 8
Patient ID: 9
Patient ID: 10
Patient ID: 11
Patient ID: 12
Patient ID: 13
Patient ID: 14
Patient ID: 15
Patient ID: 16
Patient ID: 17
Patient ID: 18
Patient ID: 19
Patient ID: 20
Patient ID: 21
Patient ID: 22
Patient ID: 23
Patient ID: 24
Patient ID: 25
Patient ID: 26
Patient ID: 27
Patient ID: 28
Patient ID: 29
Patient ID: 31
Patient ID: 32
Patient ID: 33
Patient ID: 34
Patient ID: 35
Patient ID: 36
Patient ID: 37
Patient ID: 38
Patient ID: 39
Patient ID: 40
Patient ID: 41

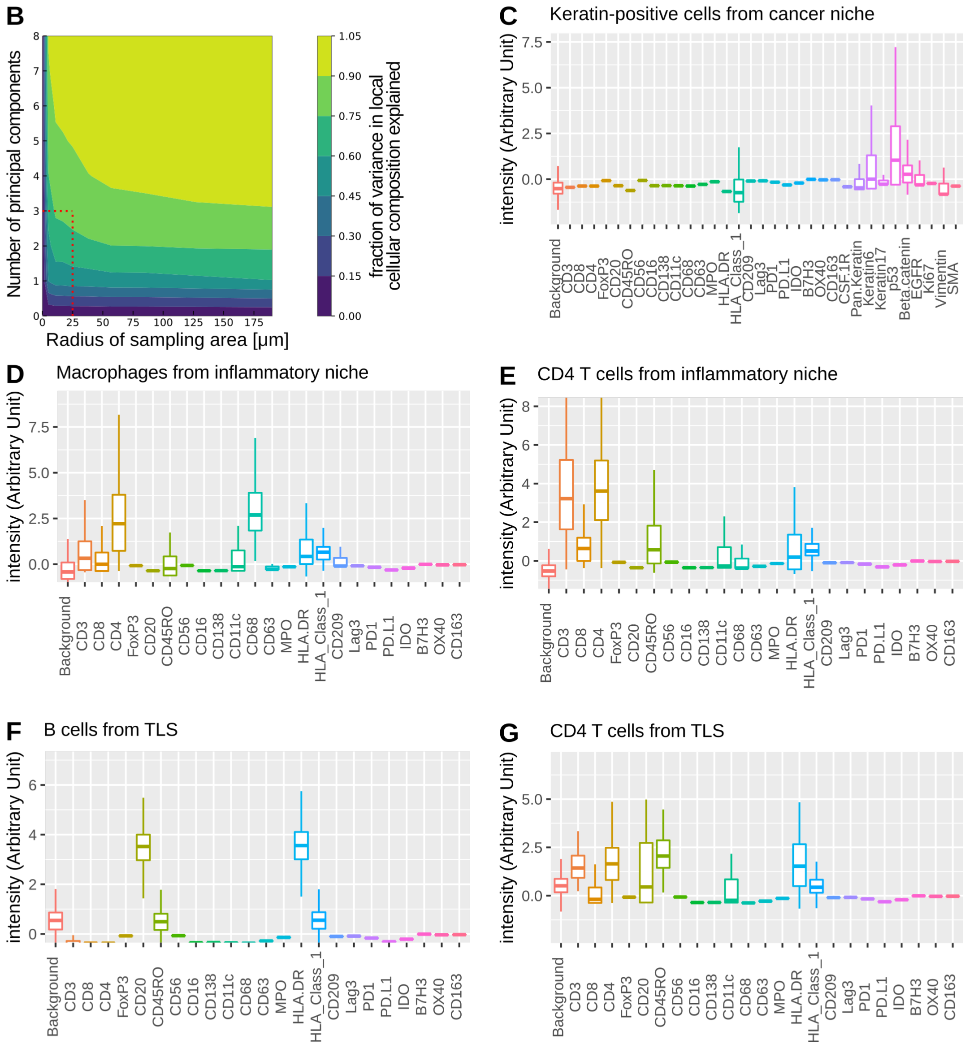


**Supplementary Figure 2: A.** Tumor architecture is difficult to interpret due to its many cell types and strong inter-sample variability. **B**. Spatial coordination of cells is detectable at a scale of 25um or larger: 82% of inter-patient variability in microscopic architecture from 4000 sampling sites of 25 micrometer radius is explained by 3 principal components. Increasing the radius beyond 25 micrometers has little impact on the amount of variance explained: this suggests that the tumor micro-environment organizes on a scale of 2-4 cells. **C-G**. Cells from the TME express different phenotypic markers depending on the TMEN in which they are found. **H.** A mix of four TMENs explain spatial variation in cellular abundance on 40 patients with TNBC. The 40 samples of Keren et al. were annotated in terms of TMENs: blue is cancer, red is inflammatory, pink is TLS, black is fibrotic/necrotic.

H

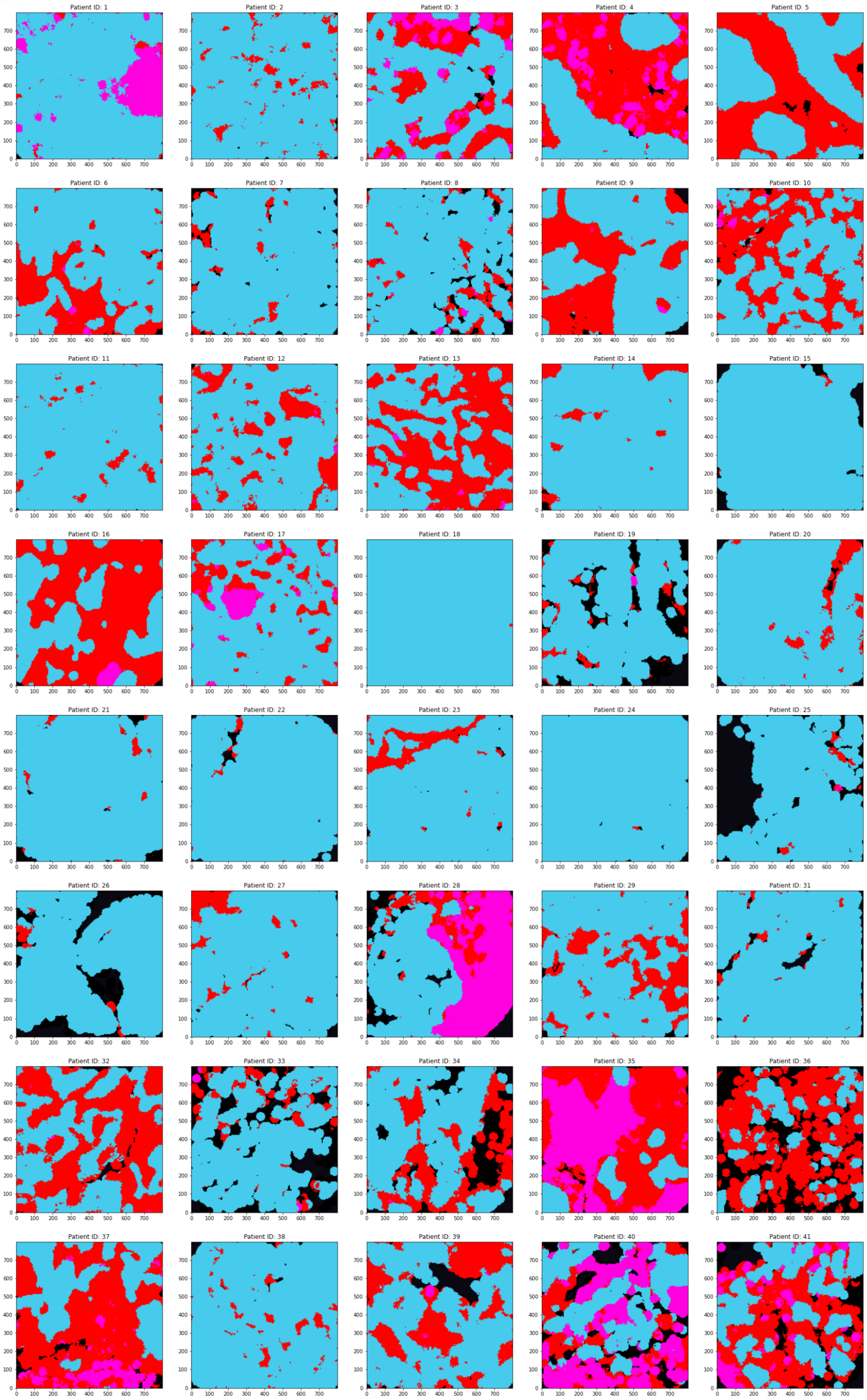
Patient ID: 1
Patient ID: 2
Patient ID: 3
Patient ID: 4
Patient ID: 5
Patient ID: 6
Patient ID: 7
Patient ID: 8
Patient ID: 9
Patient ID: 10
Patient ID: 11
Patient ID: 12
Patient ID: 13
Patient ID: 14
Patient ID: 15
Patient ID: 16
Patient ID: 17
Patient ID: 18
Patient ID: 19
Patient ID: 20
Patient ID: 21
Patient ID: 22
Patient ID: 23
Patient ID: 24
Patient ID: 25
Patient ID: 26
Patient ID: 27
Patient ID: 28
Patient ID: 29
Patient ID: 31
Patient ID: 32
Patient ID: 33
Patient ID: 34
Patient ID: 35
Patient ID: 36
Patient ID: 37
Patient ID: 38
Patient ID: 39
Patient ID: 40
Patient ID: 41
0 100 200 300 400 500 600 700

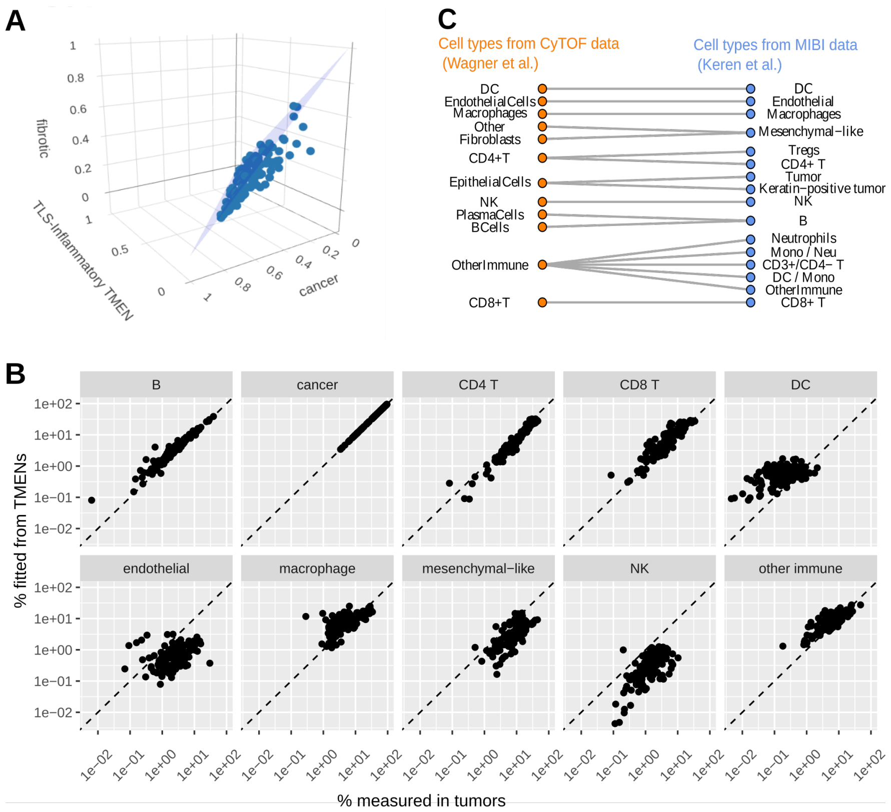


**Supplementary Figure 3: A.** In a regression analysis of macrosopic tumor cellular composition on the microscopic TMENs, the weights of microscopic TMENs fall on a unit triangle, as expected if macrosopic tumors are made of the TMENs. **B**. TMENs captures inter-patient variation in the macroscopic composition of tumors at both low and high cellular abundance. **C**. To compare the micro- and macro-architecture, cell types were mapped into a set of cell types common to MIBI (Keren et al., in blue) and CyTOF (Wagner et al., in orange) data.
